# High degree of virulence gene diversity in *Streptococcus pyogenes* isolated in Central Italy

**DOI:** 10.1101/207803

**Authors:** Daniela Bencardino, Maria Chiara Di Luca, Dezemona Petrelli, Manuela Prenna, Luca Agostino Vitali

## Abstract

Globally, *Streptococcus pyogenes* poses a continuous burden on human health, causing both self-limiting and life-threatening diseases. Therefore, studying the profile of virulence genes and their combinations is essential to monitor the epidemiology and pathogenetic potential of this important species. Thus, the aim of this study was to analyze some genetic features of clinical strains collected in Italy in 2012.

We conducted fibronectin-collagen-T antigen (FCT) region typing and *emm* typing in 122 *S. pyogenes* strains. Furthermore, several additional virulence genes were screened by polymerase chain reaction.

We found correlations between *emm* types and FCT region profiles. *emm1* strains were mainly associated with FCT2 and FCT6, while *emm89* and *emm12* strains were associated with FCT4. FCT5 was mainly represented in *emm4, emm6,* and *emm75* strains. Noteworthy, we defined subtypes for each FCT type based on the differences in single and multiple loci compared to the reference scheme used for the classification of the FCT region. In addition, new FCT types were identified. Cluster analysis based on virulence gene profiling showed a non-random distribution within each *emm* type.

This study showed the high variability of *S. pyogenes* strains and the great diversification that this pathogen has undergone during its evolution in the human host.

## Introduction

*Streptococcus pyogenes* (group A Streptococcus, GAS) is an important human pathogen that colonizes the pharynx and the skin, causing an array of diseases ranging from mild sore throat and impetigo to invasive and life-threatening infections (Cunningham, 2000).

Like most pathogens, GAS produces many virulence factors (Cunningham, 2000). Currently, eleven virulence factors have been identified as superantigens: three chromosomally encoded genes (*speG, speJ,* and *smeZ*) and eight genes harbored by temperate phages (*speA, speC, speH, speI, speK, speL, speM,* and ssa) (Friães et al., 2013). SpeB and SpeF, both mapping on the bacterial chromosome and originally described as exotoxins, are a cysteine protease and a DNAse, respectively (Friães et al., 2013). Two additional virulence factors are encoded by genes located on phages: a phospholipase A_2_ (Sla) and a DNAse (Sdn) (Friães et al., 2013). However, the primary virulence factor for GAS is the M protein (Metzgar & Zampolli, 2011). *emm* types are associated with specific superantigen profiles, and these associations differ in GAS populations collected from different geographical areas (Commons et al., 2008). It is still unknown whether environmental or biological factors may play a role in these associations.

Genes encoding several virulence factors involved in GAS pathogenesis, including the M protein, are located in the fibronectin-collagen-T antigen (FCT) region (Bessen & Kalia, 2002). Different FCT types include different subsets of *emm* types, and a genetic linkage between *emm* markers and FCT region gene products is involved in GAS tissue specificity (Kratovac et al., 2007).

Our study described 122 GAS isolated in 2012 in three regions of central Italy (Marche, Lazio, and Umbria). Our investigation focused on the distribution of virulence factors and their relationship with *emm* types as well as with FCT types. This survey may serve as the basis for further epidemiological and evolutionary studies on GAS.

## Methods

### Sample collection and identification

In 2012, 122 GAS isolates were collected from symptomatic patients with pharyngotonsillitis from hospital clinical microbiology laboratories in Roma (Ospedale Pediatrico Bambino Gesù), Perugia (Ospedale S. Maria della Misericordia), and Macerata (Laboratorio di Analisi ASL9). Isolation from throat (n = 107), rectal (n = 1), and vaginal swabs (n = 2), skin and wound (n = 5), bronchoalveolar lavage (n = 2), and normally sterile fluids (blood and pleural fluid, n = 5) was performed. Identification of clinical GAS was confirmed in our laboratory by standard procedures such as colony morphology, β-hemolysis on blood agar, serogrouping using a latex agglutination test (Streptococcal Grouping kit; Oxoid, Basingstoke, UK), bacitracin susceptibility test (Oxoid), and, additionally, by polymerase chain reaction (PCR) detection of DNaseB (Slinger et al., 2011).

### DNA extraction and *emm* typing

DNA extraction was performed using the GenElute Bacterial Genomic DNA kit (Sigma-Aldrich, St. Louis, MO, USA). *emm* typing was performed according to the Center for Disease Control and Prevention (CDC) guidelines (https://www.cdc.gov/streplab/protocolemm-type.html). To determine the *emm* type, each sequence was compared to the reference sequences available in the CDC database.

### Virulence gene profiling

The presence of *speA*, *speB*, *speC*, *speH*, *speI*, *speK*, *speL*, *speM*, *smeZ*, *ssa*, *sdn*, and *sla* virulence genes was tested by PCR using specific primer sequences (Table S1). This set of genes is referred to as non-FCT throughout the text. Briefly, each 25 µL test tube contained 1 µg of chromosomal DNA, 10 mM Tris-HCl (pH 8.3), 50 mM KCl, 1.5 mM MgCl_2_, 200 µM deoxynucleotide triphosphates (dNTPs), 1 µM oligonucleotide primers, and 0.5 U Taq polymerase (AmpliTaq Gold; Applied Biosystems, Foster City, CA, USA). The cycling program included an initial denaturation at 95°C for 2 min, followed by 30 cycles at 95°C for 40 sec, at primer-specific annealing temperature for 40 sec, at 72°C for 1 min, and one cycle at 72°C for 5 min.

### FCT typing

PCR amplification using chromosomal DNA was performed using 13 different primer pairs targeting various virulence genes within the FCT region (Table S2).^6^ The profiles of the specific products allowed the identification of FCT types, as described by Kratovac et al (Kratovac et al., 2007).

### Data analysis and statistics

Cluster analysis was done in MEGA v6 (Tamura et al., 2013) and confirmed using Statgraphics Centurion v17.1.08 (Statpoint Technologies Inc., Warrenton, VA, USA). The latter software was used to perform the principal component analysis as well as the chi-square tests when indicated (P < 0.05 was considered significant for the rejection of the null hypothesis). Display and annotation of the phylogenetic tree were obtained with Interactive Tree Of Life (iTOL) v3 online tools (Letunic et al., 2016).

## Results

### Correlation between non-FCT virulence gene profiles and *emm* types

The distribution of *emm* types within the collected strains is reported in Fig. 1A. Six *emm* types (1, 4, 89, 6, 12, and 29) accounted for nearly 70% of the population.

**Figure 1.**
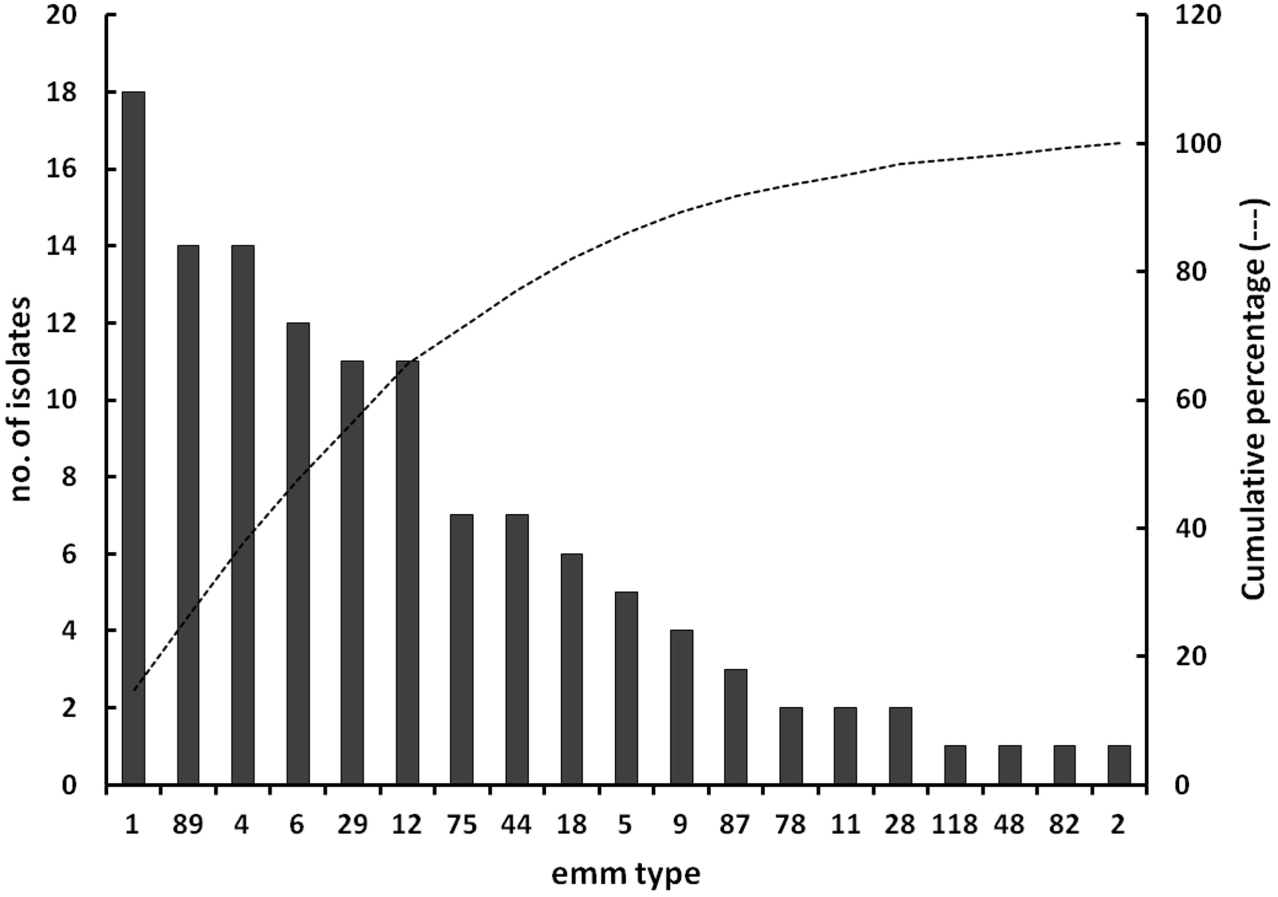
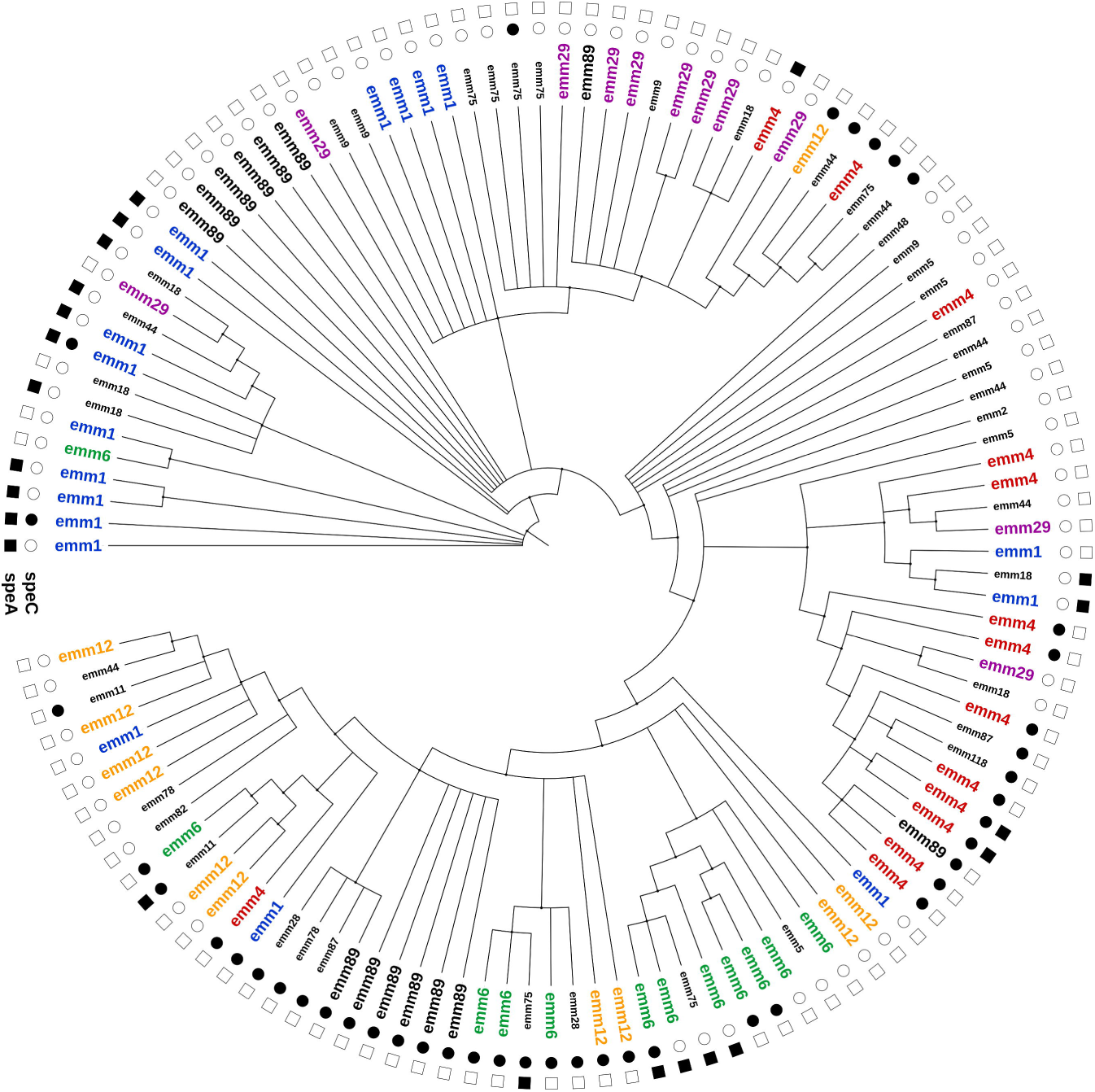
Distribution of *emm* types within the *Streptococcus pyogenes* population under investigation (A) and cluster analysis considering the whole set of virulence genes (B), showing the association with the *emm* type (inner circle) and the distribution of the genes encoding the two major superantigens, *speA* and *speC* (outer circle).

Although we detected a consistent heterogeneity of non-FCT virulence genes across all *emm* types (Fig. 1B), we considered their distribution in the most prevalent *emm* types, namely *emm1, emm4, emm89, emm6, emm12,* and *emm29* (Table 1 and Fig. 1B - labels with a larger font size). The distribution of virulence genes in each *emm* type was variable (Fig. 1B). For instance, *emm1* strains were present in 5 distant clusters as well as *emm89* strains (4 different clusters). Only 6 strains were negative to the whole set of virulence genes. *speB* was the most prevalent virulence gene present in 93 strains (76%), whereas *speK* and *sla* were the least common (15% and 14%, respectively).

**Table 1.** Distribution of virulence genes within each *emm* type group.

|  | **Total** | ***emm1*** | ***emm4*** | ***emm89*** | ***emm6*** | ***emm12*** | ***emm29*** |
| --- | --- | --- | --- | --- | --- | --- | --- |
|  | **n = 80** | **n = 18** | **n = 14** | **n = 14** | **n = 12** | **n = 11** | **n = 11** |
| *speC* | 30 | 3 | 10 | 7 | 7 | 3 | 0 |
| *speA* | 15 | 9 | 3 | 0 | 3 | 0 | 0 |
| *ssa* | 16 | 2 | 10 | 2 | 0 | 1 | 1 |
| *sdn* | 18 | 2 | 8 | 0 | 4 | 1 | 3 |
| *sla* | 11 | 0 | 1 | 0 | 10 | 0 | 0 |
| *speK* | 12 | 1 | 0 | 0 | 7 | 4 | 0 |
| *speH* | 22 | 2 | 2 | 0 | 11 | 7 | 0 |
| *speI* | 14 | 3 | 1 | 0 | 4 | 6 | 0 |
| *speL* | 14 | 0 | 2 | 1 | 0 | 2 | 9 |
| *speM* | 10 | 2 | 2 | 0 | 0 | 2 | 4 |
| *speB* | 62 | 12 | 9 | 14 | 10 | 9 | 8 |
| *smeZ* | 22 | 8 | 9 | 1 | 1 | 0 | 3 |

*speC* was identified in 71% of the *emm4,* 58% of the *emm6,* 50% of *emm89,* 27% of *emm12,* and 17% of the *emm1* strains. Strains positive to *speA* belonged to *emm1* (50%), *emm6* (25%), and *emm4* (21%). The *emm29* strains were the sole type uniformly negative to both *speC* and *speA.*

In spite of being a gene of the GAS core genome, *speB* was not detected in all strains. It was found in the *emm1* (67%), *emm4* (64%), *emm6* (83%), *emm12* (81%), *emm29* (73%), and *emm89* (100%) strains. *sdn* was found in the *emm4* (57%), *emm6* (33%), and *emm1* (11%) strains. *smeZ* was detected in the *emm4* (64%), *emm1* (44%), *emm89* (7%), and *emm6* (8%) strains. We did not observe any *emm89* strain positive to *speA, sdn, sla, speK, speH, speI,* and *speM.* We did not detect *ssa, speL,* and *speM* in the *emm6* strains as well as *sla* and *speL* in the *emm1* strains, whereas in the *emm4* strains only *speK* was not detected (Table 1).

Within *emm1, emm4, emm6,* and *emm89,* the number of different non-FCT virulence gene profiles was limited (Table S3). The analysis identified 40 genotypes, and the two most common genotypes, G38 and G39, were exclusively found in *emm89* strains (5 and 6 strains, respectively). Similarly, the 4 G7 strains belonged to *emm1.* Only genotypes G5 and G26 contained 3 strains. G5 was associated with *emm1,* whereas G26 was associated with *emm4* and *emm89* strains. The rest of genotypes were characterized by one or at least two strains and were not considered for further comparative analysis and discussion.

Variability in non-FCT virulence genes among *emm1* strains unmasked a substantial heterogeneity wherein none of the sorted principal components accounted for most of the variability (Fig. 2). There were strains containing no virulence genes and groups having *speC* or *sdn* as the major contributors to variability. Almost the same trend was found in *emm4, emm29,* and *emm89* strains. The latter group, however, was composed by strains mostly devoid of virulence genes. The *emm6* strains were characterized by the presence of *speH/speI* and *speK/sla.* Similarly, the *emm12* strains were diversified by the presence of *speC.*

**Figure 2.**
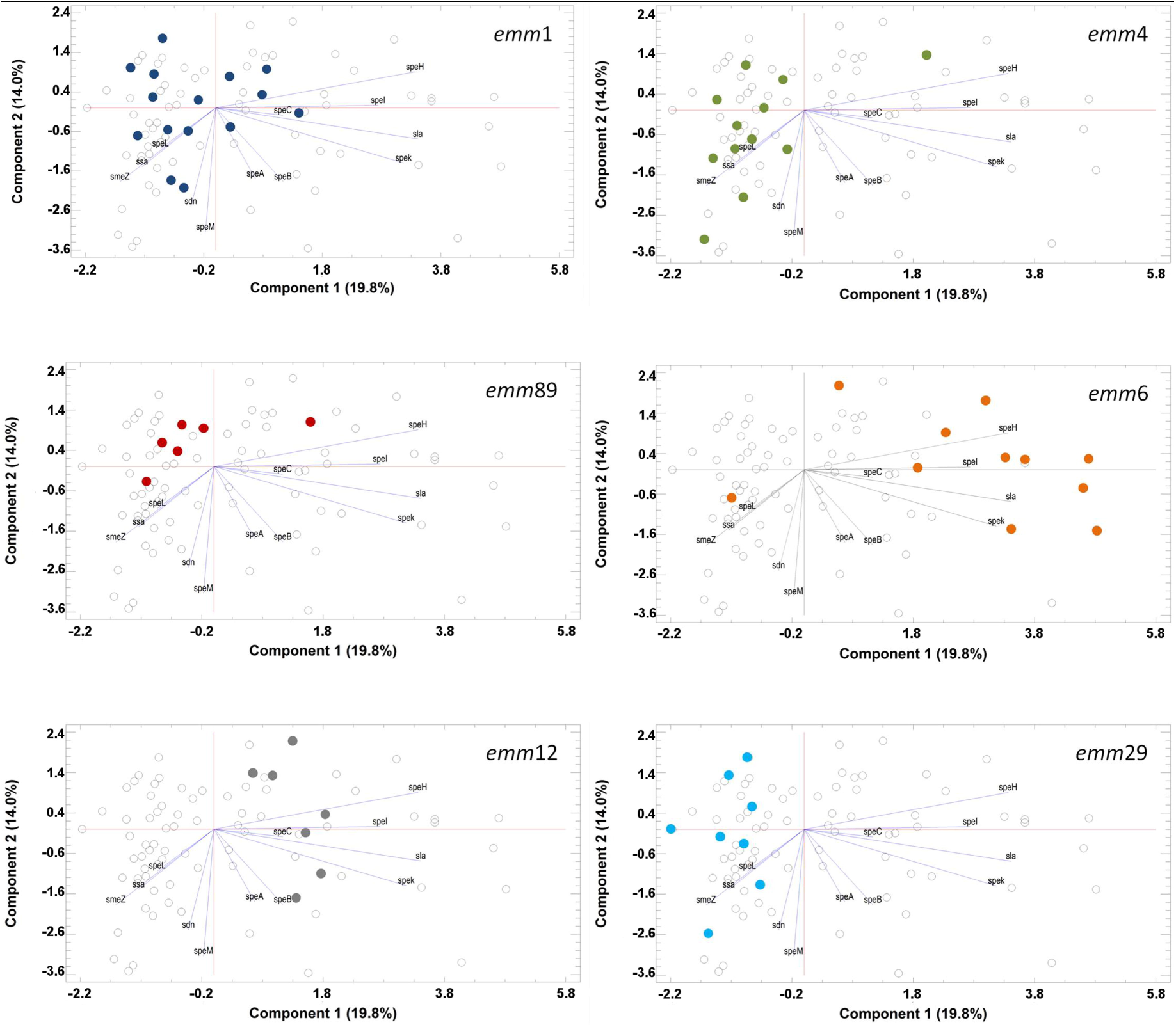
Principal component analysis (PCA) of the contribution of virulence genes to diversity. Each panel shows the localization of strains with the same *emm* type within the common 2D plot from the PCA. Each spot may result from the superimposition of more strains having the same genetic profile.

### FCT typing

A total of 41 (34%) GAS strains showed an identical structure to FCT regions described by Kratovac et al [6]. The two most frequently identified FCT regions were the FCT5 (16%) and FCT4 (10%). We constructed a classification scheme labeling with the capital letter A the FCT type matching the genetic profile described by Kratovac et al (Kratovac et al., 2007), with the capital letter B the FCT subtype differing from the reference by one gene (e.g. 5B; bridge region positive), and with the capital letter C those subtypes differing from the reference by two genes. FCT4B was the most prevalent subtype (n = 18). Compared to the reference FCT4, FCT4B lacked *fctA* (n = 12) or the bridge region (n = 6). Overall, 73 out of 122 strains (60%) showed a subtype FCT region structure not matching those already described. The association of 11 strains (9%) to a specific FCT type or subtype was not possible, because their FCT region genetic profile showed extensive differences compared to the reference.

In order to establish a possible association between the *emm* type and FCT region, we cross-tabulated *emm* types and FCT types. We run the test to determine whether or not to reject the hypothesis that the *emm* type and FCT type classifications were independent (chi-square test). Since the P-value was less than 0.001, we rejected the hypothesis that *emm* types and FCT types were independent at the 99.9% confidence level. Thus, we looked for the associations between FCT types and the most represented *emm* types. As it is shown in Fig. 3, *emm1* strains were mainly associated with FCT regions 2 and 6, while in *emm89* and *emm12* strains the most represented region was FCT4. A good proportion of both *emm4* and *emm6* strains were positive to FCT5. At last, *emm29* strains were almost completely characterized by the FCT3.

**Figure 3.**
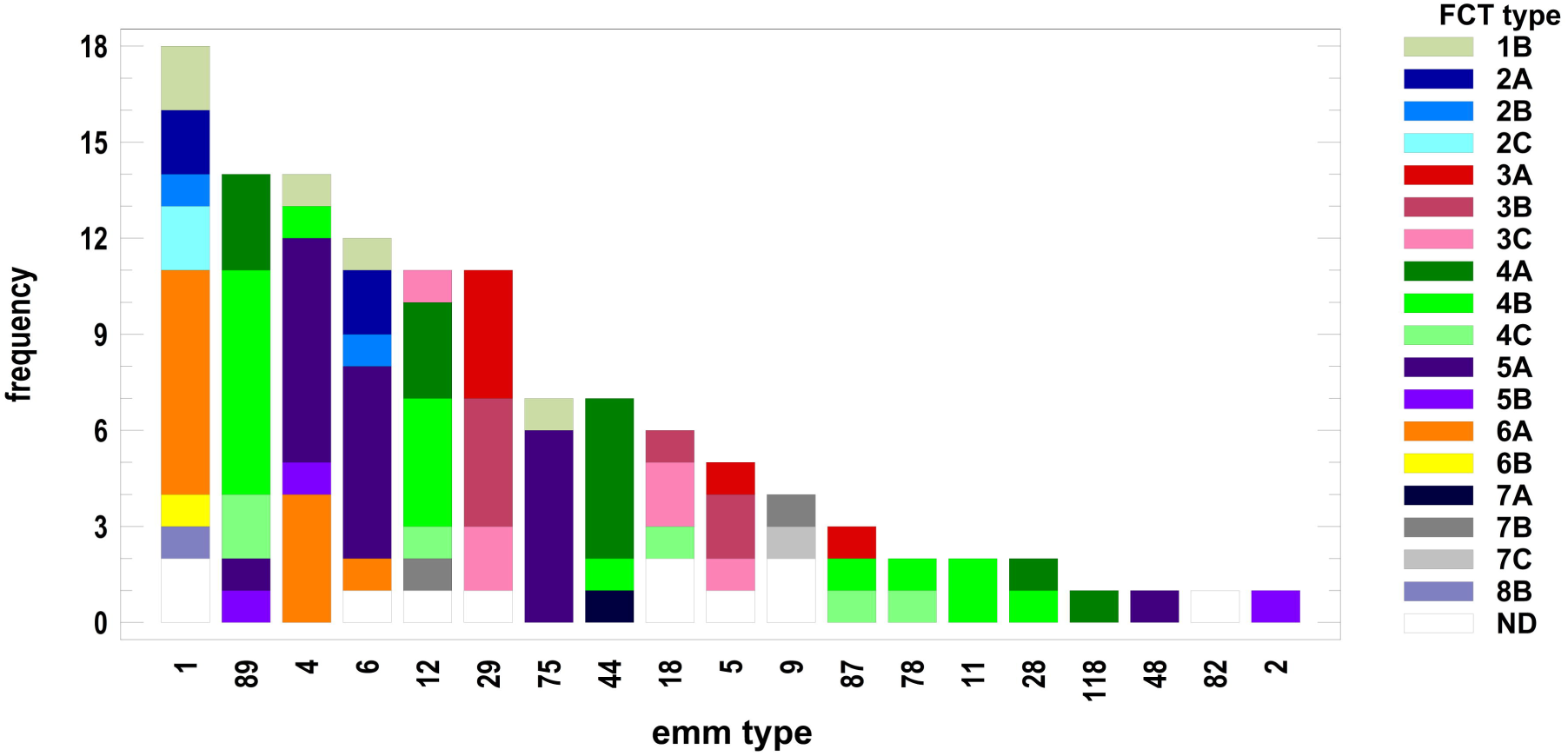
Distribution of fibronectin-collagen-T antigen (FCT) region types and subtypes within each *emm* type group of strains. Frequency on the Y-axis refers to the number of strains.

The principal component analysis based on the *emm* type, non-FCT virulence genes, and FCT region pattern showed an association between *emm* typing, FCT typing, and virulence gene profiling (Fig. 4). Component 1 was mostly and equally contributed by *fctB* and *prtF2,* followed by *fctA, nra, srtB, speB,* and *speL. rofA, prtF1,* and *srtB,* and *speH/I* contributed to variability within component 2, followed by *speK/sla, speC,* and *speB* (the latter three genes with almost the same scores). At last, component 3 included *ssa, rofA, prtF1, smeZ,* and *speC,* followed by *srtB.*

**Figure 4.**
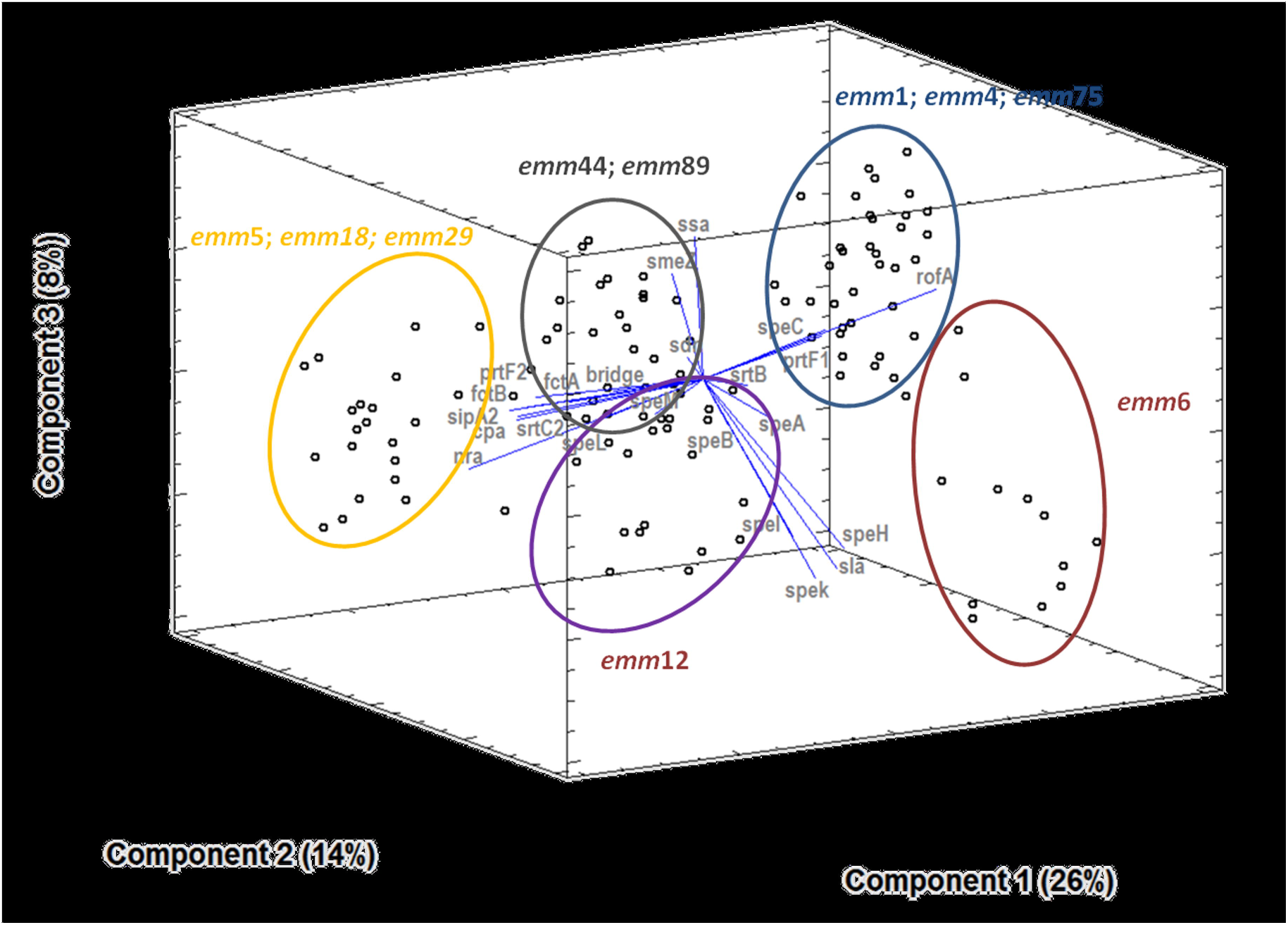
Principal component analysis (3D plot) showing the relative contribution of each virulence gene to the diversity among strains. Clustering of strains (spots) according to their *emm* type is indicated by circles of different colors.

When all virulence genes were considered (i.e. *emm,* non-FCT and FCT genes), eighty-three different virulence gene profiles were discriminated (data not shown), supporting the extremely high genetic heterogeneity of the strains.

### Geographical distribution of *emm* and FCT types

The strains were isolated from three different areas of Central Italy, namely Roma (RM; n=48), Perugia (PG; n=50) and Macerata (MC; n=26). We investigated if there was any association between the isolation area and specific virulence traits. Cross-tabulation of *emm* types by areas and the test for independence by chi-square test showed that the hypothesis of independence could be rejected *(p* < 0.001). The measurement of the degree of association between *emm* types and areas showed that there was a 43.1% reduction in error when the *emm* type was used to predict the geographical area where the strain came from. A focus on the six most common *emm* types (1, 4, 89, 6, 12, and 29) showed that the proportions of *emm1* and *emm89* were almost equally represented within the RM, PG, and MC groups. Actually, the RM group had slightly more *emm1* strains than the other two groups, whereas *emm4* strains were scarcely represented in the PG cohort. Conversely, the PG area consistently contributed to the total number of *emm6* and *emm29* strains.

The same analysis was performed to investigate the association between the area of isolation and FCT typing. The hypothesis of independence for these two variables could be rejected (*p* < 0.01). Given the overall association between the major *emm* types and the FCT types and between *emm* types and geographical area of isolation, a general association between FCT types and geographical area of isolation was expected. In fact, we observed an association between FCT3 and the PG area as well as between FCT2/FCT6 and the RM area.

## Discussion

Our investigation revealed a high variability of *emm* types and virulence genes among GAS strains from three different areas of Central Italy. Some virulence traits are mostly represented in one area over the other two. This result was not biased by the possible occurrence of major clones circulating in a single area, for instance as a result of epidemics. In fact, the large number of virulence traits considered had a high discriminatory power sorting out 83 different profiles. When compared to the total number of strains (n = 122), this number excluded clonality from being a factor influencing clustering and interpretation of variable associations.

Our data confirmed a trend observed in other countries in which *emm1, emm4, emm89,* and *emm6* (listed in descendent order) were among the prevalent circulating types (Zampaloni et al., 2003; Commons et al., 2008; Steer et al., 2009; Shea et al., 2011), followed by *emm12* and *emm29*. The association between *emm* type and source of isolation was not investigated because the number of strains isolated from throat was highly predominant and would have undermined the significance of the correspondence analysis. Genetic changes in *emm89* strains have recently generated clonal lineages responsible for invasive infections in different parts of the globe (Steer, Danchin & Carapetis, 2007; Beres et al., 2016). The high prevalence of non–invasive *emm89* type strains in our collection confirmed the increasing expansion of this *emm* type together with its propensity to switch to more virulent variants, which is of concern and claims the continuous monitoring of its diffusion into the population (Latronico et al., 2016). The analysis of the recorded patterns of virulence genes revealed the presence of major clones as already described in previous studies (Vlaminckx et al., 2003; Commons et al., 2008; Friães et al., 2012). We found *emm4* strains with the G25 pattern of virulence genes. They had a wide distribution and were recorded in invasive and non-invasive infections over a long period of time (Commons et al., 2008; Friães et al., 2012). Among the *emm6* strains, the FCT5 and G35 pattern of virulence genes were identified. A similar pattern was present in invasive GAS isolated in The Netherlands from 1992 and 1996 (Vlaminckx et al., 2003). In addition, we described *emm1*/G16 pattern strains that differed for the presence of *smeZ* from *emm1* strains isolated in Portugal (Friães et al., 2012).

This distribution of *emm* types revealed a complex epidemiological scenario of GAS strains. The detection of 12 virulence genes showed a great degree of clonal heterogeneity within the GAS collection.

We observed a great structural variability within the FCT region, higher than that previously described (Kratovac et al., 2007). New FCT type variants were recorded, indicating an extremely high degree of genetic shuffling among different parts of the FCT region in GAS. This is most probably related to the evolutionary pressure of the immune system on the different elements exposed on the surface of GAS and encoded by genes mapping within the FCT region such as fibronectin-binding proteins and pili (Mora et al., 2005). Further and continuous molecular epidemiologic studies are needed to increase our understanding of possible associations of virulence determinants and their variants that facilitate host-pathogen interactions.

## Acknowledgements

This work was supported by funds from the Italian Ministry of Education, University, and Research (MIUR, grant FUTURO IN RICERCA number RBFR10X4YN_001 to D.P.).

